# DeNoFo: a file format and toolkit for standardised, comparable de novo gene annotation

**DOI:** 10.1101/2025.03.31.644673

**Authors:** Elias Dohmen, Margaux Aubel, Lars A. Eicholt, Paul Roginski, Victor Luria, Amir Karger, Anna Grandchamp

**Affiliations:** Institute for Evolution and Biodiversity, University of Münster, Hüfferstrasse 1, 48151, Münster, Germany; Institute for Integrative Biology of the Cell (I2BC), Université Paris-Saclay, CEA, CNRS, 1 Av. de la Terrasse Bâtiment 21, 91198, Gif-sur-Yvette, France; Department of Neuroscience, Yale School of Medicine, New Haven, 06510, CT, USA; Department of Systems Biology, Harvard Medical School, Boston, 02115, MA, USA; Division of Genetics and Genomics, Boston Children’s Hospital, Harvard Medical School, Boston, 02115, MA, USA; IT-Research Computing, Harvard Medical School, Boston, 02115, MA, USA; Aix Marseille University, INSERM, TAGC, UMR S1090,Marseille, France

**Keywords:** de novo genes, proto-genes, annotation format, standardisation, comparability

## Abstract

**Motivation:** *De novo* genes emerge from previously non-coding regions of the genome, challenging the traditional view that new genes primarily arise through duplication and adaptation of existing ones. Characterised by their rapid evolution and their novel structural properties or functional roles, *de novo* genes represent a young area of research. Therefore, the field currently lacks established standards and methodologies, leading to inconsistent terminology and challenges in comparing and reproducing results.

**Results:** This work presents a standardised annotation format to document the methodology of *de novo* gene datasets in a reproducible way. We developed DeNoFo, a toolkit to provide easy access to this format that simplifies annotation of datasets and facilitates comparison across studies. Unifying the different protocols and methods in one standardised format, while providing integration into established file formats, such as fasta or gff, ensures comparability of studies and advances new insights in this rapidly evolving field.

**Availability and Implementation:** DeNoFo is available through the official Python Package Index (PyPI) and at https://github.com/EDohmen/denofo. All tools have a graphical user interface and a command line interface. The toolkit is implemented in Python3, available for all major platforms and installable with pip and uv.

## Introduction

*De novo* genes are defined as genes that emerge from previously non-genic sequences within a genome. Since the early 2000s, significant progress has been made in understanding the mechanisms underlying their emergence (Long et al., 2003; Begun et al., 2007; Cai et al., 2008; Knowles and McLysaght, 2009; Tautz and Domazet-Lo?so, 2011; Carvunis et al., 2012; Wissler et al., 2013; Schmitz et al., 2018; Van Oss and Carvunis, 2019; Heames et al., 2020; Zheng and Zhao, 2022; Parikh et al., 2022; Vakirlis et al., 2022; Zhao et al., 2024; Rich and Carvunis, 2023; Iyengar and Bornberg-Bauer, 2023; Aldrovandi et al., 2024; Peng and Zhao, 2024). The prevalence of *de novo* genes across all domains of life is now well-documented, and their biological roles within respective species remain an active area of research (Li et al., 2010; Gubala et al., 2017; Baalsrud et al., 2018; Zhang et al., 2019; Rivard et al., 2021; Lange et al., 2021; Broeils et al., 2023; Vakirlis and Kupczok, 2024).

A major challenge in the study of *de novo* genes is the accurate discrimination of recently originated *de novo* genes from those that have emerged via alternative mechanisms (Schmitz et al., 2018). This challenge stems from the scarcity of automated pipelines capable of automatically detecting and annotating *de novo* genes, which have only recently been developed (Arendsee et al., 2019; Casola et al., 2022; Roginski et al., 2024). Consequently, the majority of *de novo* genes have been identified through the implementation of non-standardized pipelines that integrate diverse software and custom scripts. Moreover, the characterisation criteria of *de novo* emergence, which varies across studies (Van Oss and Carvunis, 2019), is a crucial factor in the identification of *de novo* genes. Past definitions have varied, incorporating different evolutionary stages and criteria (Keeling et al., 2019; Weisman, 2022), thresholds for how much of a gene must have arisen *de novo* to qualify as such (McLysaght and Hurst, 2016), criteria for characterising the absence of homology (Casola, 2018; Vakirlis et al., 2020; Weisman et al., 2022), and different modes of emergence (Pereira et al., 2024). The ambiguities in defining and identifying *de novo* gene emergence have prompted the usage of diverse methodologies. While exploring various methods is essential for a relatively young field of research, the lack of consensus on definitions and the use of varying methodologies, have hindered the effective comparison of results across studies (Schmitz et al., 2018; Aubel et al., 2023; Grandchamp et al., 2025). As a consequence, datasets comprising *de novo* genes from the same species can differ considerably, and in some cases, may not even be overlapping, depending on the methodologies and definitions employed to determine the *de novo* status of a given gene. Several studies investigating *de novo* gene emergence in the same species show large discrepancies. For example, in Roginski et al. (2024), the authors detected 89 *de novo* genes in humans, while Vakirlis et al. (2022) identified 155 de novo Open Reading Frames (ORFs), and Dowling et al. (2020) identified 2,749 human-specific *de novo* ORFs. The observed discrepancies can be attributed to methodological differences. Specifically, Roginski et al. (2024) detected *de novo* genes through the analysis of annotated genomes, Vakirlis et al. (2022) utilized ORFs identified via ribosome profiling as candidates, and Dowling et al. (2020) sought candidates within transcriptomic data.

Similarly, Heames et al. (2020) detected 66 *de novo* genes in *Drosophila melanogaster*, while Roginski et al. (2024) detected 92 and Peng and Zhao (2024) 555 *de novo* genes. In the same species Zheng and Zhao (2022) identified 993 *de novo* ORFs and a fifth study detected an average of 1,548 *de novo* ORFs (Grandchamp et al., 2023). These discrepancies primarily stem from variations in methodology and underlying data sources.

These methodological discrepancies and different definitions are discussed in more detail in Grandchamp et al. (2025), along with the categorisation and standardisation of these methods for the annotation format presented here. The characterisation and understanding of *de novo* gene emergence is a young research field, and for future advancements, the exploration of different methodologies remains crucial. However, the wide variety of methodologies complicates the comparability of datasets and hinders progress.

Consequently, there is an essential need for a standardised and comparable approach to ensure reproducibility and facilitate the comparison of results across studies without restrictions on the applied methods and definitions. To address this need, we present DeNoFo, a toolkit designed to automate and streamline the annotation process for *de novo* gene detection and validation as well as rapid and straightforward comparison of methodology across studies.

## Tool Description

### The standardised annotation format

The present diversity in definitions, terms, protocols, tools and pipelines mentioned above to investigate *de novo* gene evolution impedes comparability of datasets and studies.

While it is important to enable a certain flexibility regarding the methodology to detect and describe *de novo* genes, other factors such as the meaning of terms or comparable description of approaches should be standardised to enable reproducibility and comparability. We propose the following annotation format to categorise and standardise the methods used for *de novo* gene detection while still allowing for the required flexibility to include a great variety of approaches. With this, information on used protocols and tools from both bioinformatic and wet lab approaches can be integrated into widely-accepted file formats in genomics.

The here described format focuses on the methodology used for the detection of *de novo* genes in different studies, in contrast to providing information about individual genes. By omitting gene-centric properties and only covering methodological aspects, all genes of a study can be covered by a single description of their methodology and studies become comparable between each other.

Based on the standardised and categorised methods for bioinformatic detection of *de novo* genes we compiled (Grandchamp et al., 2025), the format is structured into six main sections (see Fig. 1): input data, evolutionary information, homology filter, non-coding homologs, lab verification and hyperlinks(URLs/DOIs).

**Fig. 1:**
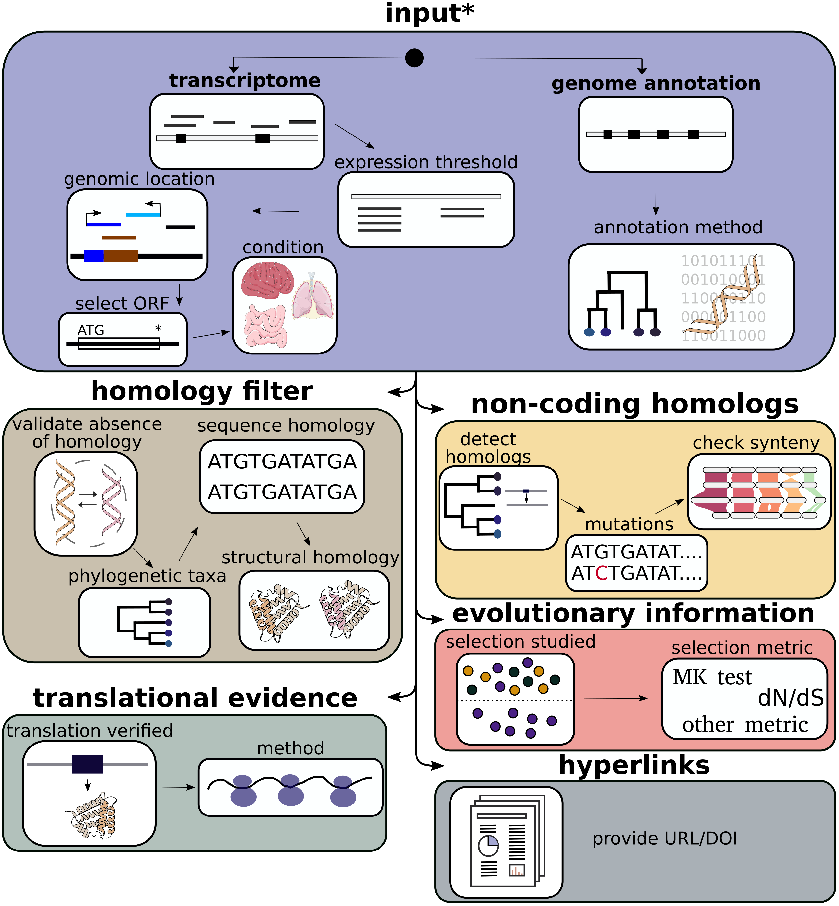
Structure of the de novo gene annotation format with six main sections and related subsections. The star indicates non-optional sections. This figure has been designed using resources from https://bioicons.com/.

Our annotation format with the here described information can be stored either in a separate file and shared as such or can be included in widely-accepted bioinformatic file formats. The default DeNoFo file format is JSON-based and human-readable to a high degree (*.dngf file extension). To allow for integration into widely-accepted file types and formats, such as fasta or gff, we developed additionally a short string encoding of the DeNoFo format, which can be added to the fasta header or the additional info column in gff files. This short string encoding of the DeNoFo format is not human-readable due to its compressed information content. For easy use and transfer of information into this proposed unified format as well as extraction of information from the mentioned file types annotated with this format, we developed a toolkit to facilitate the usage.

### Toolkit Functionality

The tools of the DeNoFo toolkit assist in the conversion of workflows into the required format, the automated annotation of *de novo* genes with the format, and the extraction of information in an easily accessible and interpretable manner. Additionally, a tool is provided for the comparison of two such *de novo* gene set annotations with regard to their similarities and differences in methodological approaches. All tools presented here are available with a graphical user interface (GUI) and a command-line interface (CLI).

The **DeNoFo-Questionnaire** represents the core component of the toolkit, serving as a guide that directs users through the required sections of the format via interactive queries. These queries offer either pre-populated options or the option to enter custom answers. To ensure user convenience, the user is permitted to move between questions and modify previous answers as required. The resulting DeNoFo annotation is saved to a user-specified file in a JSON-like *de novo* gene file format (dngf, that is also the file extension).

The **DeNoFo-Converter** is a tool designed for the purpose of converting the DeNoFo format from one file type to another. To illustrate this function, one may consider the process of annotating sequences in a fasta or gff file with an annotation file in the dngf format that has already been saved. To enhance user convenience, the tool offers various options for annotating or extracting only annotations from a selection of sequences.

The **DeNoFo-Comparator** is a tool designed for the purpose of comparing two studies or datasets that have been annotated in the DeNoFo format. This tool facilitates the loading of two different DeNoFo annotations and the generation of a report that either highlights the similarities or differences between the two studies or datasets. The report is presented in an easily interpretable, human-readable formatting.

The help function and tooltips of the GUI and CLI versions of each tool, as well as the DeNoFo user manual, contain further explanations and examples with regard to the options and functionality of the toolkit and its constituent tools.

## Implementation

The DeNoFo toolkit has been developed in modern Python (version 3.10 or higher) and is released under the GPL v3 license. All GUI versions are implemented using PyQt6 (Limited, 2024). The DeNoFo format is implemented as a set of pydantic models (Colvin et al., 2025) which contain the definitions and validations for the DeNoFo format.

The pydantic models are independent of the tools with specific user interfaces. Decoupling the models from the tools allows anyone to interact with the format programmatically without using our tools, while ensuring a high level of robustness of the format. For example, any *de novo* gene detection software could implement its own routine to automatically generate a valid annotation format file, so that a user does not need to manually generate one using the DeNoFo-Questionnaire.

For easy and fast setup the DeNoFo toolkit supports installation via pip or uv and has been tested on all major platforms (Linux, Windows, MacOS).

## Conclusion

The DeNoFo format and toolkit presented here provide the community with an easily accessible, unified and comparable standard for methodology annotation for *de novo* genes. The overarching objective of this initiative is to enhance the comparability, interpretability and reproducibility of data sets and findings within this new and heterogeneous field of research.

The format has been designed to be as flexible and open as possible to accommodate the diverse methodological approaches in this field, while ensuring a unified annotation and reporting with a focus on best practice. To this end, the format has been designed to allow for changes and adaptation as this field of research moves forward. In order to facilitate this, the format has been implemented in such a way that additions and changes can be made with as few compatibility issues as possible. This approach enables the incorporation of new databases or software for future studies in the format, without requiring researchers to update their existing annotations.

The community is encouraged to contribute to the format by adding new approaches, databases, software and more, as well as requesting or contributing features and functionality that would improve the accessibility, usability or comparability of the format, toolkit and datasets.

In conclusion, the unified format and toolkit for *de novo* gene annotation, developed with and for the community, will hopefully pave the way for faster progress and deeper understanding in this field of research, while being continuously adapted and improved in conjunction with innovative methods and novel tools to advance the field of *de novo* gene evolution.

## Supporting information

SupplementaryMaterial-Case-Studies-DENOFO

## Author contributions statement

AG was responsible for the conceptualisation of the format with contributions from all authors. ED was responsible for the software development and implementation of the toolkit, with feedback and testing from all authors, particularly AG, MA and LE. AG and ED jointly handled project administration. MA and AG were responsible for the visualisation of the format as a figure for the paper and designed the layout and colour scheme of the toolkit. VL and AK contributed to developing the evidence criteria and the questionnaire for evaluating de novo gene status. The original draft of the manuscript was written by ED and AG, with subsequent editing and review of the first version by MA and LE. The final manuscript was edited and reviewed by all authors. Erich Bornberg-Bauer provided general administrative support and acquisition of the financial support for the project leading to this publication.

## Acknowledgments

We thank Erich Bornberg-Bauer (EBB), Nikolaos Vakirlis, Li Zhao, Anne Lopes, Bharat Ravi Iyengar and Andreas Lange for their useful feedback during the planning phase of the project. ED was funded by the Deutsche Forschungsgemeinschaft (DFG, German Research Foundation) – 503348080. This grant with the additional grant number BO2544-22-1 was awarded to EBB. MA received funding from the Volkswagen foundation with grant code 98183 awarded to EBB. AG was supported by the Deutsche Forschungsgemeinschaft priority program “Genomic Basis of Evolutionary Innovations” (SPP 2349) BO 2544/20-1 to EBB, and by the (HFSP) Postdoctoral Fellowship (Grant No. 2023-981550) awarded to EBB, Anne-Ruxandra Carvunis and Christine Brun. LAE has been supported by EMBO Scientific Exchange Grant 10944. VL was supported by NIH grant R01NS095654 (to Nenad Sestan).

For Figure 1 the following icons were taken from https://bioicons.com/: brain-1 icon (colour changed), healthy-lung icon and intestine icon by Servier https://smart.servier.com/ licensed under CC-BY 3.0 simple DNA backbone icon (colour changed) by Marnie-Maddock https://github.com/ MarnieMaddock licensed under CC-BY 4.0 Riparian plots icon by Chenxin-Li https://github.com/cxli233 licensed under CC-BY4.0 Protein monochrome icon (colour changed) by DBCLS https://togotv.dbcls.jp/en/pics.html licensed under CC-BY 4.0

