## SupplementaryMaterial-Case-Studies-DENOFO for "DeNoFo: a file format and toolkit for standardised, comparable de novo gene annotation": Human_differences_Vakirlis-2022_vs_Dowling-2020.txt

Differences between Vakirlis et al. (2022) and Dowling et al. (2020):

	inputTranscriptome:

		expressionLevel:

			differing values in Vakirlis et al. (2022) and Dowling et al. (2020):
			0.1
			0.5

		transContextChoice:

			values in Vakirlis et al. (2022) but not in Dowling et al. (2020):
			custom choice

			values in Dowling et al. (2020) but not in Vakirlis et al. (2022):
			overlap gene
			intronic
			intergenic
			antisense

		customGeneticContext:

			values in Vakirlis et al. (2022) but not in Dowling et al. (2020):
			previously annotated non-coding transcripts 
			upstream of coding ORFs on coding transcripts
			downstream of coding ORFs on coding transcripts 
			on transcripts lacking coding ORFs but which belong to a transcript family with a member annotated as coding

		transORFChoice:

			values in Vakirlis et al. (2022) but not in Dowling et al. (2020):
			custom choice

			values in Dowling et al. (2020) but not in Vakirlis et al. (2022):
			longest ORF

		customORF:

			values in Vakirlis et al. (2022) but not in Dowling et al. (2020):
			ORF-RATER score >=0.8

		transcriptomeInfo:

			differing values in Vakirlis et al. (2022) and Dowling et al. (2020):
			human induced pluripotent stem cells (iPSCs), iPSC-derived cardiomyocytes, human foreskin fibroblasts (HFFs), and HFFs infected with cytomegalovirus
			brain, cerebellum, heart, lung, kidney, and testes 

DeNovoGeneAnnotation:

	evolutionaryInformation:

		values in Vakirlis et al. (2022) but not in Dowling et al. (2020):
		selection: codeml, FEL

	phylogeneticTaxa:

		taxSpecificity:

			differing values in Vakirlis et al. (2022) and Dowling et al. (2020):
			lineage-specific
			species-specific

		taxonID:

			taxID:

				differing values in Vakirlis et al. (2022) and Dowling et al. (2020):
				human-vertebrata
				9606

	homologyFilter:

		seqType:

			values in Vakirlis et al. (2022) but not in Dowling et al. (2020):
			RNA

		threshold:

			values in Vakirlis et al. (2022) but not in Dowling et al. (2020):
			coverage [%]
			50.0

			differing values in Vakirlis et al. (2022) and Dowling et al. (2020):
			e-value
			0.0001, 0.001

		dataBase:

			values in Vakirlis et al. (2022) but not in Dowling et al. (2020):
			RefSeq

			values in Dowling et al. (2020) but not in Vakirlis et al. (2022):
			custom choice

		customDB:

			values in Dowling et al. (2020) but not in Vakirlis et al. (2022):
			transcriptomes of Pan paniscus, Pan troglodytes, Gorilla gorilla, Pongo pygmaeus, Macaca mulatta, Mus musculus

	nonCodingHomologs:

		values in Vakirlis et al. (2022) but not in Dowling et al. (2020):
		enablingMutations: True
		synteny: 
			anchors: genome alignment

	translationalEvidence:

		translationEvidence:

			values in Dowling et al. (2020) but not in Vakirlis et al. (2022):
			mass spectrometry

		customTranslationEvidence:

			values in Vakirlis et al. (2022) but not in Dowling et al. (2020):
			hexamer score

			values in Dowling et al. (2020) but not in Vakirlis et al. (2022):
			hexamer scores

		studyURL:

			values in Vakirlis et al. (2022) but not in Dowling et al. (2020):
			https://doi.org/10.1016/j.celrep.2022.111808

			values in Dowling et al. (2020) but not in Vakirlis et al. (2022):
			https://doi.org/10.1093/gbe/evaa194
