## SupplementaryMaterial-Case-Studies-DENOFO for "DeNoFo: a file format and toolkit for standardised, comparable de novo gene annotation": Human_differences_Vakirlis-2022_vs_Roginski-2024.txt

Differences between Vakirlis et al. (2022) and Roginski et al. (2024):

DeNovoGeneAnnotation:

	inputData:

		values in Vakirlis et al. (2022) but not in Roginski et al. (2024):
		transcriptome

		values in Roginski et al. (2024) but not in Vakirlis et al. (2022):
		annotated genome

	inputAnnotGenome:

		values in Roginski et al. (2024) but not in Vakirlis et al. (2022):
		annotGenomeChoice: homology-based approach

	inputTranscriptome:

		values in Vakirlis et al. (2022) but not in Roginski et al. (2024):
		expressionLevel: 0.1
		transContextChoice: custom choice
		customGeneticContext: 
			previously annotated non-coding transcripts 
			upstream of coding ORFs on coding transcripts
			downstream of coding ORFs on coding transcripts 
			on transcripts lacking coding ORFs but which belong to a transcript family with a member annotated as coding
		transORFChoice: custom choice
		customORF: ORF-RATER score >=0.8
		transcriptomeInfo: human induced pluripotent stem cells (iPSCs), iPSC-derived cardiomyocytes, human foreskin fibroblasts (HFFs), and HFFs infected with cytomegalovirus

	evolutionaryInformation:

		selection:

			differing values in Vakirlis et al. (2022) and Roginski et al. (2024):
			codeml, FEL
			codeml

		taxonID:

			taxID:

				differing values in Vakirlis et al. (2022) and Roginski et al. (2024):
				human-vertebrata
				human-hominoidea

	homologyFilter:

		seqType:

			values in Vakirlis et al. (2022) but not in Roginski et al. (2024):
			RNA

		threshold:

			values in Vakirlis et al. (2022) but not in Roginski et al. (2024):
			coverage [%]
			50.0

		dataBase:

			values in Vakirlis et al. (2022) but not in Roginski et al. (2024):
			RefSeq

			values in Roginski et al. (2024) but not in Vakirlis et al. (2022):
			NCBI NR

	nonCodingHomologs:

		enablingMutations:

			values in Vakirlis et al. (2022) but not in Roginski et al. (2024):
			True

		synteny:

			anchors:

				values in Vakirlis et al. (2022) but not in Roginski et al. (2024):
				genome alignment

				values in Roginski et al. (2024) but not in Vakirlis et al. (2022):
				gene anchors

			softwareSyntenySearch:

				values in Roginski et al. (2024) but not in Vakirlis et al. (2022):
				DENSE

	translationalEvidence:

		values in Vakirlis et al. (2022) but not in Roginski et al. (2024):
		translationEvidence: 
			ribosome profiling
			custom choice
		customTranslationEvidence: hexamer score

	studyURL:

		values in Vakirlis et al. (2022) but not in Roginski et al. (2024):
		https://doi.org/10.1016/j.celrep.2022.111808

		values in Roginski et al. (2024) but not in Vakirlis et al. (2022):
		https://doi.org/10.1093/gbe/evae159
