## SupplementaryMaterial-Case-Studies-DENOFO for "DeNoFo: a file format and toolkit for standardised, comparable de novo gene annotation": Two_case_studies_denofo.pdf

### Two Case Studies on de novo Genes

#### Human and fly *de novo* genetic elements

To demonstrate the capabilities of our newly developed toolkit DENOFO, we analysed (i) three different studies that identified *de novo* genetic elements (dngs) in humans and (ii) three other studies that identified dngs in flies. Between the studies of the same taxon, we compared the number of detected dngs, differences and similarities in methodology, using our newly developed DENOFO annotation format and toolkit.

We annotated these six studies in our developed standardised DENOFO annotation format through the DENOFO toolkit and compared them, providing qualitative insights into differences and similarities of dngs studies.

We provide the annotation files and all detailed pairwise comparisons between the three studies regarding both the differences and the similarities in the Supplementary Material "case_study_human_dngs" and "case_study_fly_dngs".

#### Case study 1: De novo genetic elements in human

The amount of detected dngs between the three selected studies for human dngs (see Fig. 1) differs by orders of magnitude. These differences in reported numbers of dngs is likely due to different methods, databases and thresholds used. To assess those discrepancies, we annotated the studies with our DENOFO toolkit and applied the pairwise comparator to each study combination (see Tab. 1).

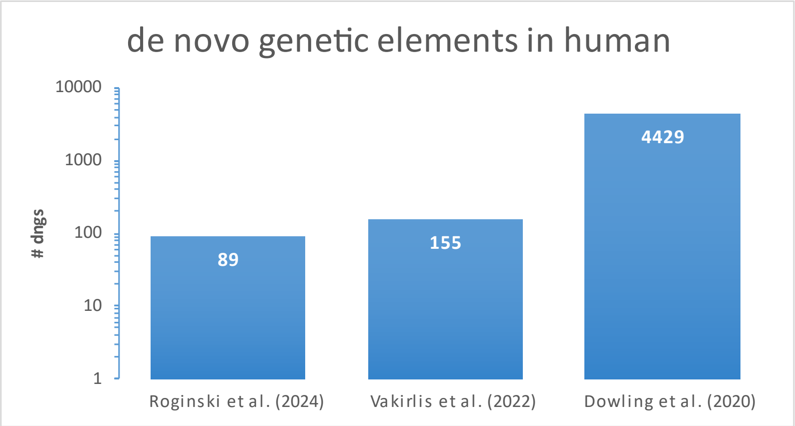


*Figure 1: Number of dngs detected in Homo sapiens by three different studies.*

*Table 1: Similarities and differences between studies on human dngs*

| | Roginski et al. (2024) | Vakirlis et al. (2022) | Dowling et al. (2020) |
|---|---|---|---|
| **Input** | genome | transcriptome | transcriptome |
| **ORF choice** | | custom | longest |
| **Specificity** | lineage | lineage | species |
| **Homology filter** | protein sequences | protein sequences | protein sequences |
| **E-value** | 0.0001 | 0.0001 | 0.001 |
| **Database** | NCBI nr | RefSeq | custom |
| **Synteny** | gene anchors | whole genome alignment | |
| **Translational evidence** | | ribosome profiling | ribosome profiling |
| **Selection** | codeml | codeml | |

The two studies with the biggest difference in number of detected dngs, Roginski et al. (2024) and Dowling et al. (2020), share only one methodological overlap in total across all reported standardised features. The studies only overlap in their use of protein sequences for homology filtering (see "Human_similarities_Dowling-2020_vs_Roginski-2024.txt" in Supplementary Material, representing the output of the denofo-comparator).

The two studies that are closest by number of detected dngs, Roginski et al. 2024 and Vakirlis et al. 2022, overlap in two annotations. Next to the already above-mentioned use of protein sequences for homology filtering, it is the reporting of lineage-specific dngs (in contrast to species-specific or population-specific ones). Although the studies by Vakirlis et al. and Dowling et al. show a large discrepancy in number of detected dngs (155 vs. 4429), their methodologies overlap, such as the used input data being in both cases the human transcriptome (in contrast to the genome) and the application of ribosome profiling for evidence of translation.

The only two overlaps in methodology between these studies, could already explain the discrepancy in the number of detected dngs. However, it becomes apparent, that even if the same type of input data (transcriptome) and the same method for translational evidence (ribosome profiling) were used, the number of detected dngs in humans still differs by orders of magnitude between those studies.

Dowling et al. (2020) and Roginski et al. (2024) are farthest apart regarding the number of detected dngs (see Supplementary Material "Human_differences_Dowling-2020_vs_Roginski-2024.txt", representing the output of the denofo-comparator). The most apparent difference is the used input data, which is a human transcriptome in Dowling et al. (2020), but an annotated genome in Roginski et al. (2024). While Roginski et al. (2024) identify lineage-specific dngs, Dowling et al. focuses on strictly species-specific genes, leading to a difference in number of detected dngs.

Also, selected thresholds are easy to identify here as a source of differences in number of detected dngs: Roginski et al. selected a way stricter e-value threshold of 0.0001 for homology filtering than Dowling et al. with 0.001 as an e-value threshold. The stricter e-value threshold results in a lower number of considered dngs, which fits to the much lower number of identified dngs.

Apart from only methodological differences leading to discrepancies in numbers of identified dngs, the annotation and comparison through DENOFO allows to extract useful information contained in the studies, which might be relevant for readers. As an example, we can see in the differences from the denofo-comparator output that Roginski et al. report additional evolutionary information in the form of selection studied through codeml.

#### Case study 2: De novo genetic elements in fruit fly

The number of detected dngs between the three selected studies on fly dngs (see Fig. 2) vary greatly. This discrepancy most likely stems from different methodologies applied, used databases and selected thresholds. To better asses the similarities and differences, we annotated the studies with our DENOFO toolkit and applied the pairwise comparator to each study combination (see Tab. 2).

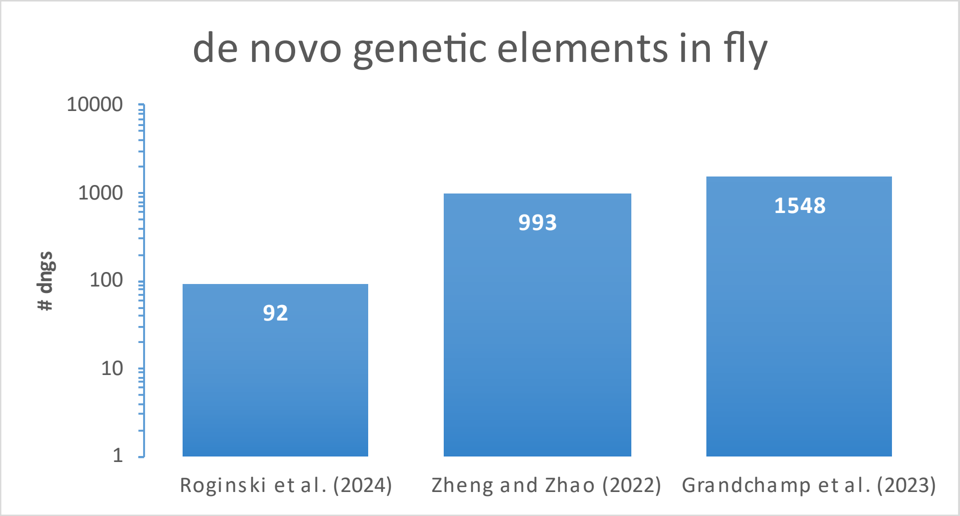


*Figure 2: Number of dngs detected in Drosophila melanogaster by three different studies.*

*Table 2: Similarities and differences between studies on flydngs*

| | Roginski et al. (2024) | Zheng & Zhao (2022) | Grandchamp et al. (2023) |
|---|---|---|---|
| Input | genome | transcriptome | transcriptome |
| ORF location | | intergenic, antisense, intronic overlapping | intergenic, antisense, intronic overlapping |
| Specificity | lineage | lineage | species |
| Homology filter | protein sequences | protein sequences | protein sequences |
| E-value | 0.0001 | | 0.1 |
| Database | NCBI nr | custom | custom |
| Synteny | gene anchors | | gene anchors |
| Translational evidence | | ribosome profiling, MS | |
| Selection | codeml | | dN/dS |
| Enabling mutations | | | yes |

Roginski et al. (2024) and Grandchamp et al. (2023), the two studies with the biggest difference in number of detected dngs in flies, share only two methodological similarities in total across all reported standardised features. These similarities are the use of protein sequences for homology filtering and using gene anchors for synteny detection (see “"Fly_similarities_Roginski-2024_vs_Grandchamp-2023.txt” in Supplementary Material, representing the output of the denofo-comparator). The two studies that are closest by number of detected dngs, Zheng & Zhao (2022) and Grandchamp et al. (2023), overlap in type of input data (transcriptome), which transcripts are considered (intergenic, antisense, intronic overlapping), protein sequences for homology filtering. Hence, the discrepancy in the number of detected dngs, must stem from the data analysis like the use of ribosome profiling and mass spectrometry (MS) for translational evidence in Zheng & Zhao (2022).

It is both difficult and time consuming to compare and assess the datasets and methodologies of published data on detected dgns manually. The standardised annotation with the developed DENOFO toolkit provides an overview of overlaps between datasets in a convenient way. Next to such surprising similarities in methodology, DENOFO also allows to compare the differences between the studies, which can help us to identified why studies using the same input data come to different conclusions.

To analyse the differences in more detail, we focus here on the two studies by Roginski et al. (2024) and Grandchamp et al. (2023) that are farthest apart regarding the number of detected dngs (see Supplementary Material “Fly_differences_Roginski-2024_vs_Grandchamp-2023.txt”, representing the output of the denofo-comparator).

We identify differences in the input data, which is a transcriptome in Grandchamp et al. (2023), but an annotated genome in Roginski et al. (2024). While Roginski et al. identify lineage-specific dngs, Grandchamp et al.'s are strictly species-specific. This alone can explain already a large difference in number of detected dngs. Also, selected thresholds are easy to identify here as a source of differences in number of detected dngs: Roginski et al. selected a way stricter e-value threshold of 0.0001 for homology filtering than Grandchamp et al. with 0.1 as an e-value threshold. The stricter e-value threshold will result in a lower number of dngs filtered out, which contrasts with the much lower number of identified dngs. Homology filtering was based on a custom database of Drosophila and Dipteran proteomes in Grandchamp et al., while Roginski et al. Used the NCBI nr database. The database of more closely related species in Grandchamp et al. can have led to the higher amount of dngs, which were filtered out in Roginski et al. due to matches in more distantly related species. Apart from only methodological differences leading to discrepancies in numbers of identified dngs, the annotation and comparison through DENOFO allows to extract useful information contained in the studies, which might be relevant for readers. As an example, we can see in the differences from the denofo-comparator output that Grandchamp et al. report information about enabling mutations, which is missing in Roginski et al. However, Roginski et al. report additional evolutionary information in the form of selection studied through codeml, while Grandchamp et al. report selection information based on dN/dS values.

#### Conclusions

The numbers of detected dngs between the selected studies on both, human and fly dngs (see Fig. 1-2, note logarithmic scale), differ by orders of magnitude. This large discrepancy might stem from different applied methodologies, databases and selected thresholds. Yet, it can be difficult to assess and compare which exact methods were applied and where they differ. With the information provided through the standardised annotation format and analysed and processed for easy comparison by the DENOFO tools, we can gain insights into the specifics of differences and similarities between these studies. Additionally, we learn about the impact of specific methodological differences the more studies are annotated this way and can be compared and analysed in a feasible way.
